# Green Elegance T2T: An upgraded telomere-to-telomere genome assembly and annotation of looseleaf lettuce (*Lactuca sativa* var. *crispa*)

**DOI:** 10.1101/2024.11.20.624396

**Authors:** Bin Zhang, Xue Liu, Haifeng Ding, Yesheng Yang, Jinfu Tang, Dayong Li

## Abstract

Lettuce (*Lactuca sativa* L.), a globally important vegetable crop, is cultivated in various horticultural varieties. Looseleaf lettuce is one of the main varieties that are consumed. Here, we report a telomere-to-telomere (T2T) genome assembly of looseleaf lettuce (*L. sativa* var. *crispa* cv. Green Elegance). By combining the sequencing data of previously reported Green Elegance v1.2, the 42.20 Gb clean ultra-long reads produced by Oxford Nanopore Technology were used to close the gaps and upgrade the genome assembly. After filling 11 gaps in Green Elegance v1.2, the final gapless telomere-to-telomere genome assembly is 2.58 Gb in length with a contig N50 of 282.47 Mb, containing nine centromeres and 18 telomeres. The genome contained 41,375 protein-coding genes, of which 99.10% were functionally annotated. This T2T genome of looseleaf lettuce will be valuable for the identification of genetic variation and the advancement of lettuce breeding.

## Introduction

Lettuce (*Lactuca sativa* L.), originally from the Mediterranean region, is a widely consumed leafy vegetable crop with significant global economic importance, generating approximately $16.6 billion in production value worldwide in 2021 (Food and Agriculture Organization of the United Nations, 2023). Looseleaf lettuce, a significant horticultural variety, is widely consumed in salads, hamburgers, and hotpots. Given its enormous economic value, the genomic research on looseleaf lettuce is essential and could greatly aid the development of lettuce breeding programs and relevant industries.

Telomere-to-telomere (T2T) gapless genome assembly could supply novel insights into genome variation and function (Garg et al., 2024). The first lettuce genome, form the crisphead lettuce type (*L. sativa* var. *capitata* cv. Salinas), was sequenced and published in 2017 (Reyes-Chin-Wo et al., 2017). Owing to progress in sequencing approaches, several other lettuce genomes have been sequenced in recent years. In 2023, two wild relatives (*L. saligna* and *L. virosa*) (Xiong et al. 2023a; Xiong et al. 2023b) and one stem lettuce (*L. sativa* var. *angustana* cv. Yanling1) were released (Shen et al., 2023). In 2024, the first chromosome-level genome assembly of looseleaf lettuce (*L. sativa* var. *crispa* cv. Green Elegance v1.2) (Zhang et al., 2024), a gapless genome assembly of cutting lettuce (*L. sativa* var. *cutting* cv. Black Seeded Simpson) (Cao et al., 2024), and the first T2T genome of Roman lettuce (*L. sativa* var. *roman* cv. PKU06) were published (Wang et al., 2024). Until now, only one T2T genome assembly of Roman lettuce has been released (Wang et al., 2024). More T2T genome assemblies of other cultivated lettuce varieties would provide molecular insights into variation and domestication within the Lactuca genus.

In this study, we present the first gapless T2T genome assembly of looseleaf lettuce, *L. sativa* var. *crispa* cv. Green Elegance T2T, produced using ONT (Oxford Nanopore Technology) ultra-long sequencing reads, based on *L. sativa* var. *crispa* cv. Green Elegance v1.2. The Green Elegance T2T genome represents a high-quality assembly, superior in accuracy compared to other previously released lettuce genomes. This genomic resource will facilitate genome comparison within the Lactuca genus and aid in the characterization of genome structure.

## Results and discussion

### T2T assembly of *Lactuca sativa* var. *crispa* cv. Green Elegance

To obtain the T2T genome of *L. sativa* var. *crispa* cv. Green Elegance, we performed ONT ultra-long sequencing (Figure 1A and Table S1). A total of 86.89 Gb of raw data was generated, and 42.20 Gb of clean ONT ultra-long reads were obtained with a clean N50 length of 100,242 bp after filtering adapters and low-quality reads (Table 1 and Table S1). Combining the HiFi, ONT, and Hi-C sequencing data from the genome of Green Elegance v 1.2 (Zhang et al., 2024), a gapless assembly of the genome of 2.58 Gb was obtained with a contig N50 of 282.47 Mb after filling 11 gaps in the genome of Green Elegance v 1.2 genome (Figure 1B, Table 1, and Table S1). Following the anchoring contigs, we ultimately obtained the first telomere-to-telomere level genome assembly of looseleaf lettuce (Green Elegance T2T) with 18 telomeres, nine centromeres, and no gaps on nine chromosomes (Figure 1C, Figure 1D, Table 1 and 2). The lengths of the telomeres ranged from 10.29 to 71.31 Kb (Table 2). Overall, there are no apparent structural assembly errors as indicated in the Hi-C interaction map, and Green Elegance T2T achieved a quality value of 46.07, an LAI value of 30.36, and a BUSCO score of 98.51%, the highest among previously released lettuce genomes, validating the integrity and accuracy of the assembly (Figure 1B and Table 3).

**Table 1.** Comparison of genome assemblies of Green Elegance T2T, Green Elegance (V1.2), Salinas (V11), Yanling1 (SL1.0), PKU06 (LsT2T), Black Seeded Simpson (CutV01)

| Feature | Green Elegance T2T | Green Elegance (V1.2) | Salinas (V11) | Yanling1 (SL1.0) | PKU06 (LsT2T) | Black Seeded Simpson (CutV01) |
| --- | --- | --- | --- | --- | --- | --- |
| Contig N50 | 282.47 Mb | 205.47 Mb | 12.52 Mb | 4.71 Mb | 320.6 Mb | 320 Mb |
| Longest contig | 369.98 Mb | 282.47 Mb | 75.54 Mb | 20.98 Mb | 409.41 Mb | 408.43 Mb |
| Total size of assembled contigs | 2.58 Gb | 2.59 Gb | 2.59 Gb | 2.60 Gb | 2.59 Gb | 2.58 Gb |
| TE proportion | 77.23% | 77.11% | 74.2% | 81.08% | 75.76% | 71.88% |
| Centromeres annotated | 9 | 9 | 0 | 0 | 9 | 9 |
| Telomeres annotated | 18 | 14 | 16 | 5 | 18 | 17 |
| Gap number | 0 | 11 | 384 | 1,404 | 0 | 1 |

**Table 2.** Statistics of Green Elegance T2T chromosomes.

| Group | Group Length (bp) | Telomere length (Upstream/Downstream, Kb) | Centromere Start (bp) | Centromere End (bp) | Centromere Length (bp) |
| --- | --- | --- | --- | --- | --- |
| Chr 1 | 257,076,624 | 20.83/12.61 | 180,010,191 | 186,348,708 | 6,338,517 |
| Chr 2 | 236,300,515 | 23.83/29.47 | 85,582,847 | 90,316,231 | 4,733,384 |
| Chr 3 | 320,751,947 | 13.05/28.85 | 228,007,447 | 230,598,831 | 2,591,384 |
| Chr 4 | 407,152,990 | 13.16/10.40 | 273,271,592 | 277,099,720 | 3,828,128 |
| Chr 5 | 369,975,375 | 11.32/18.37 | 115,760,824 | 121,515,957 | 5,755,133 |
| Chr 6 | 205,473,523 | 20.21/14.89 | 112,357,708 | 115,098,823 | 2,741,115 |
| Chr 7 | 213,801,038 | 15.48/13.03 | 115,746,959 | 120,568,250 | 4,821,291 |
| Chr 8 | 342,781,751 | 14.51/71.31 | 262,494,994 | 265,808,459 | 3,313,465 |
| Chr 9 | 229,447,819 | 10.29/ 17.89 | 131,163,399 | 134,946,656 | 3,783,257 |

**Table 3.** Quality values, BUSCO and LAI assessments of Green Elegance T2T, Green Elegance (V1.2), Salinas (V11), Yanling1 (SL1.0), PKU06 (LsT2T), Black Seeded Simpson (CutV01)

| Type | Green Elegance T2T | Green Elegance (V1.2) | Salinas (V11) | Yanling1 (SL1.0) | PKU06 (LsT2T) | Black Seeded Simpson (CutV01) |
| --- | --- | --- | --- | --- | --- | --- |
| Complete BUSCOs (C) | 1590 (98.51%) | 1588 (98.39%) | 1589 (98.45%) | 1586 (98.27%) | 1575 (97.60%) | 1579 (97.81%) |
| Complete and single-copy BUSCOs (S) | 1554 (96.28%) | 1551 (96.10%) | 1554 (96.28%) | 1535 (95.11%) | 1557 (96.5%) | —— |
| Complete and duplicated BUSCOs (D) | 36 (2.23%) | 37 (2.29%) | 35 (2.17%) | 51 (3.16%) | 35 (2.2%) | —— |
| Fragmented BUSCOs (F) | 6 (0.37%) | 8 (0.50%) | 3 (0.19%) | 8 (0.50%) | 4 (0.20%) | —— |
| Missing BUSCOs (M) | 18 (1.12%) | 18 (1.12%) | 22 (1.36%) | 20 (1.24%) | 18 (1.1%) | —— |
| Total Lineage BUSCOs | 1,614 | 1,614 | 1,614 | 1,614 | 1,614 | 1,614 |
| LAI | 30.36 | 17.34 | 18.13 | 12.01 | —— | —— |
| QV | 46.07 | 44.45 | 28.65 | 42.38 | 58.03 | —— |

**Figure 1.**
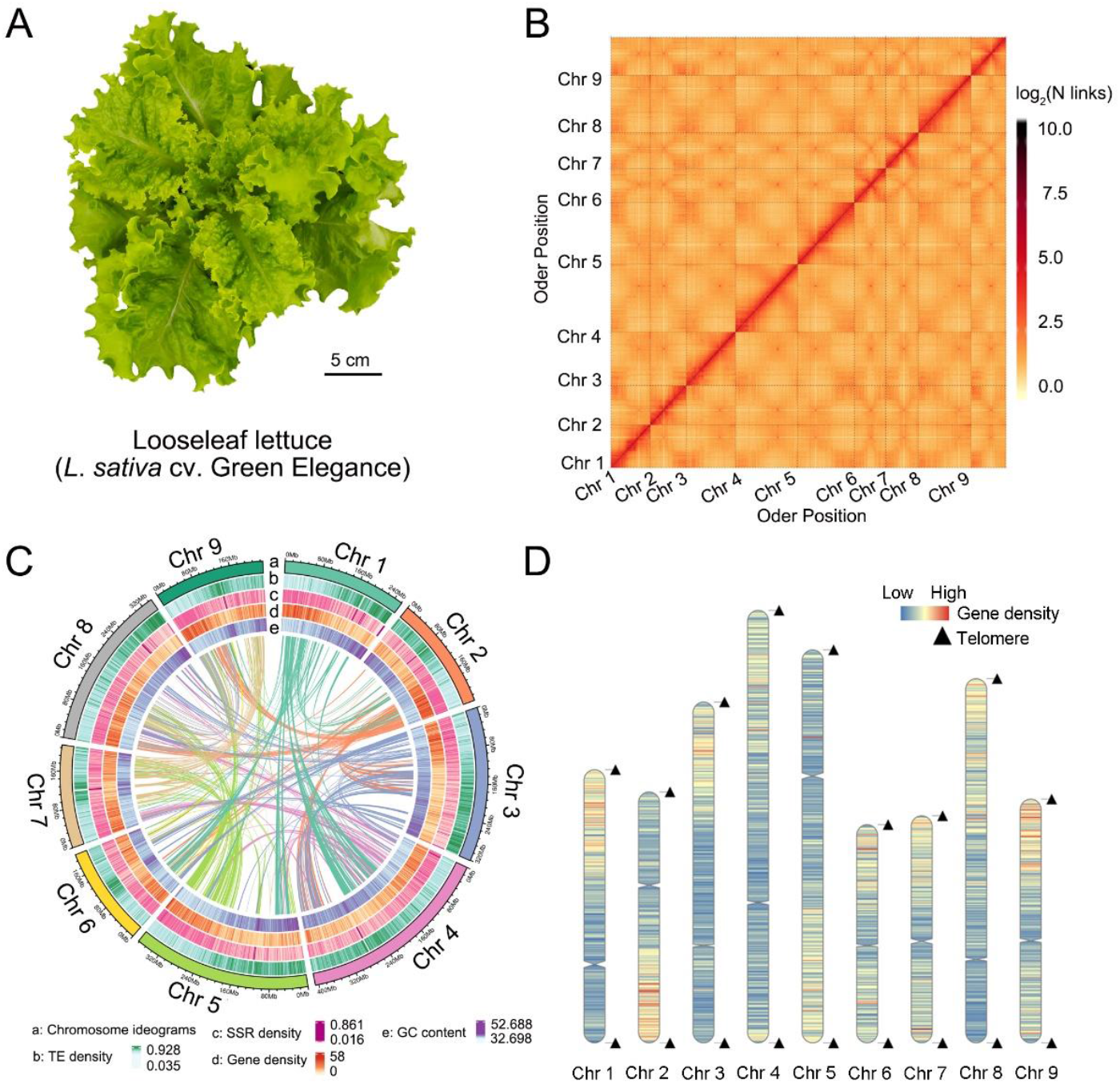
T2T Genome assembly of the looseleaf lettuce (*L. sativa* var. *crispa* cv. Green Elegance) A. Looseleaf lettuce (cv. Green Elegance) at 30-day of growing. B. Hi-C heatmap of Green Elegance T2T genome. The intra-chromosome interaction density is displayed in different colours and calculated using a bin size of 800 K. C. Circos plot of genome assembly of Green Elegance T2T. Chromosome ideograms (a), TE density (b), SSR density (c), gene density (d), and GC content (e) were exhibited from the outer to the inner layer, respectively. D. The distribution of centromeres and telomeres on chromosomes of Green Elegance T2T. Eighteen telomeres, nine centromeres and gene density (bin size = 1 Mb) were shown on chromosomes.

### Genome annotation and gene prediction

Approximately 2.00 Gb of transposon elements (77.23% of the genome) and 106.62 Mb of tandem repeats (4.13% of the genome) were identified (Table S2). A total of 45,770 protein-coding genes were predicted in the Green Elegance genome, with an average length of 4,019 bp. Of these, 99.12% (45,365 genes) could be functionally annotated with known genes, conserved domains, and Gene Ontology terms (Table S3). This gene number is consistent with that observed in two other T2T lettuce genomes (*L. sativa* var. *roman* cv. PKU06 and Green Elegance T2T), though the functional annotation rate in Green Elegance (T2T) is notably higher (99.12%) than in PKU06 (Table S3).

### Detection of centromeres and telomeres

The centromeric sequences for the nine chromosomes were assembled, with an average length of 4.21 Mb. The longest centromere was 6.34 Mb on chromosome 1, while the shortest was 2.59 Mb on chromosome 3 (Table 2). Centromeric regions exhibited higher repeat sequence density, including transposable elements (TEs) and simple sequence repeats (SSRs), and lower gene density (Figure 1C, 1D).

### Genomic comparison of Green Elegance T2T with the previously released lettuce genome

The genome assembly of *L. sativa* var. *crispa* cv. Green Elegance (T2T) in this study demonstrates substantial improvements in genome continuity (contig N50), gaps, and the number of centromeres and telomeres compared to other previously released lettuce genomes, including L. sativa var. capitata cv. Salinas (V11), L. sativa var. angustana cv. Yanling1 (SL1.0), L. sativa var. roman cv. PKU06, and L. sativa var. cutting cv. Black Seeded Simpson (Cutv01) (Table 3). Additionally, the higher ratio of complete BUSCOs, Quality Value (QV), and LAI values in Green Elegance (T2T) compared to other lettuce genomes further underscores the high quality of this genome assembly (Table 1, Table 3).

## Material and methods

### Plant material

The sequenced lettuce variety in this study, looseleaf lettuce (*Lactuca sativa* var. *crispa* cv. Green Elegance), was provided by the Beijing Vegetable Research Center, Beijing Academy of Agriculture and Forestry Science, Beijing, China (Fig. 1). The growth conditions of Green Elegance were described in Zhang et al. (2024). The healthy leaves of 30-day-old seedlings of Green Elegance were collected and immediately frozen in liquid nitrogen for genome sequencing on the ONT (Oxford Nanopore Technologies) ultra-long sequencing platform (ONT ultralong-reads-2rd) (Table S1).

### Genome sequencing

ONT ultra-long sequencing was performed to obtain the Telomere-to-Telomere (T2T) genome of Green Elegance. High-quality genomic DNA was extracted using the CTAB method (cetyltrimethylammonium bromide) (Abu Almakarem et al., 2012) and fragmented to ∼8 Kb. The ONT ultra-long sequencing library was then constructed using the Ligation Sequencing Kit 1D (Nanopore, SQK-LSK109) and sequenced on the PromethION sequencer (Oxford Nanopore Technologies).

### Genome assembly

Combining 124.16 Gb high-accuracy Circular Consensus Sequencing (CCS) clean data and 32.55 Gb ONT sequencing (ONT ultralong-1st) clean data in Zhang et al. (2024), ONT ultralong reads (ONT ultralong-reads-2rd) were used to fill gaps in Green Elegance v1.2 (Table S1). Assembly of contigs was conducted using hifiasm (v 0.16) software with the default parameters (Cheng et al., 2021). For anchored contigs, the Hi-C reads were mapped to the polished Green Elegance T2T genome using BWA (bwa-0.7.17) with the default parameters. Then, scaffold assembly was performed using the agglomerative hierarchical clustering method in Lachesis (Burton et al., 2013). The Hi-C contact heat map was drawn using Hicexplorer v3.7 (Wolff et al., 2020). Gap filling was performed by quartTeT software (https://github.com/aaranyue/quarTeT) using contigs and was conducted by TGS GapCloser (1.2.0) (https://github.com/BGI-Qingdao/TGS-GapCloser) using third-generation sequencing and ultra-long ONT data. Finally, the final T2T genome was obtained.

### Genome quality assessment

The Illumina short-reads and PacBio long-reads data were mapped back to the assembly, and the alignment file was analyzed with Qualimap v.2.2.2 (Okonechnikov et al., 2016). The assembled genome was subjected to BUSCO (Benchmarking Universal Single-Copy Orthologs) v5.2.2 and CEGMA (Core Eukaryotic Genes Mapping Approach) (v2.5) to evaluate the integrity of the final genome assembly (Simão et al., 2015; Parra et al., 2007). Merqury (v1.3) (Rhie et al., 2020) was used for Quality Value (QV) accuracy assessment and the LAI (LTR Assembly Index) (Ou et al., 2018) value was used for genome assembly continuity of *L. sativa* var. *crispa* cv. Green Elegance T2T.

### Repeat element identification and gene annotation

Transposon elements (TE) were identified using a mix of homology-based and *de novo* methods, including the Dfam (v3.5) database, RepeatModeler (http://www.repeatmasker.org/RepeatModeler/) (Flynn et al., 2020), LTRharvest (v1.5.9) (Ellinghaus et al., 2008), LTR_finder (v2.8) (Xu and Wang, 2007), and LTR_retriever (Ou and Jiang, 2018). The TE sequences were classified by RepeatMasker (v4.12) (Tarailo-Graovac and Chen, 2009). Tandem repeats were annotated by Tandem Repeats Finder (TRF 409) (Benson, 1999) and MIcroSAtellite identification tool (MISA v2.1) (Beier et al., 2017).

Three approaches were integrated for gene annotation, including *de novo* prediction, homology search, and transcript-based assembly. The *de novo* gene models were predicted using Augustus (v3.1.0) (Stanke et al., 2004) and SNAP (Korf, 2004). GeMoMa (v1.7) software (Keilwagen et al., 2019) was used for homolog-based alignment using reference gene models. Hisat (v2.1.0) (Kim et al., 2015) and Stringtie (v2.1.4) (Pertea et al., 2015) were employed for the transcript-based prediction.

Gene functions were annotated by comparison with the Non-Redundant (NR), EggNOG (Huerta-Cepas et al., 2019), KOG (Koonin et al., 2004), TrEMBL (Boeckmann et al., 2003), InterPro (Mitchell et al., 2015), Swiss-Prot protein databases (Boeckmann et al., 2003), Gene Ontology (GO) databases (Ashburner et al., 2000), and the Kyoto Encyclopedia of Genes and Genomes (KEGG) database (Kanehisa et al., 2012). The protein motifs and domains were annotated using InterProScan (v5.34-73.0) (Jones et al., 2014) and PFAM databases (Finn et al., 2014).

### Identification of telomeres and centromeres

The potential telomere repeat units in the genome were predicted using TIDK (https://github.com/tolkit/telomeric-identifier). Based on the predicted repetitive units, the sequences and locations of potential telomere sequences were identified using FindTelomeres (https://github.com/JanaSperschneider/FindTelomeres).

The potential centromere repeats were predicted using Centromics (https://github.com/ShuaiNIEgithub/Centromics). Then, the predicted centromere repeats were compared to the genome to identify the positions and sequences of centromeres. The chromosome maps were drawn using the R package RIDeogram, marking telomeres, centromeres, and gene density.

## Code Availability

No custom code was used for this study. Unless otherwise specified, all data analyses were conducted using published bioinformatics software with default settings.

## Data availability

The ONT ultra-long sequencing raw data used for genome assembly in this study are available in the Genome Sequence Archive (GSA) (Chen et al., 2021b) at the National Genomics Data Center (NGDC), Beijing Institute of Genomics (China National Center for Bioinformation) (CNCB-NGDC Members and Partners, 2023), Chinese Academy of Sciences (https://bigd.big.ac.cn/gsa). The accession number is CRA020495. The genome assembly and annotation files are available in the Genome Warehouse (GWH) (Chen et al., 2021a) at NGDC (accession number is GWHFHHL00000000.1) and on Figshare (https://doi.org/10.6084/m9.figshare.27823269).

## Acknowledgments

This research was supported by the Key Project at Central Government Level: the Ability Establishment of Sustainable Use for Valuable Chinese Medicine Resources (2060302), the Beijing Innovation Consortium of Agriculture Research System (BAIC01-2023). The authors thank Dr. Tianhua He (Murdoch University, Australia) and Dr. Man Zhou (University of Maryland, USA) for their critical reading and comments on the manuscript.

## Author contributions

D.L. and B.Z. designed and coordinated the study; X.L., H.D., Y.Y., J.T., and B.Z. collected and prepared plant samples; X.L. and B.Z. performed the bioinformatic analyses; B.Z. and D.L. drafted the manuscript; J.T., B.Z., and D.L. revised the manuscript. All authors approved the final manuscript.

